# SILICA-SNC: FROM STREAMLINES TO NETWORKS

**DOI:** 10.64898/2026.09.23.753938

**Authors:** Lei Wu, Vince Calhoun

**Affiliations:** TReNDS Center Georgia State University, Georgia Institute of Technology, Emory University Atlanta, Georgia 30303

**Keywords:** diffusion MRI, tractography, SILICA, structural network coupling, independent component analysis

## Abstract

Whole brain tractography contains rich trajectory-level information about white matter organization, yet conventional structural connectomes capture only part of it. We introduce Structural Network Coupling (SNC), a network representation enabled by SILICA (Streamline Independent Component Analysis). SILICA decomposes tractography into structural source components and their streamline-specific loading profiles, while SNC characterizes relationships among these components through their shared trajectory expression. The resulting component-level network retains direct correspondence to the underlying white-matter pathways and supports network visualization and graph-based analysis. SNC produced anatomically interpretable coupling among trajectory-resolved white-matter components and revealed a highly organized group architecture that independently recapitulated commissural, projection, and association systems. It also showed strong hemispheric segregation and structured cross-system coupling, and was highly stable across split-half samples. SNC extends SILICA from trajectory decomposition to network-level organization and provides a flexible framework for future structural and multimodal network analysis.

## 1. INTRODUCTION

Tractography and structural connectomics describe the same white-matter architecture at very different scales. A tractogram is inherently trajectory-based, comprising large numbers of continuous paths with complex spatial relationships. Conventional structural connectomes, in contrast, typically summarize these trajectories as pairwise relationships between network nodes [1–3]. The nodes may be defined anatomically, through cortical parcellation, or using data-driven representations [4], while edges are commonly quantified using streamline counts or related weighting measures. This provides an important representation of anatomical connectivity, but it does not exhaust the structural information available from tractography, particularly the organization encoded in trajectory-level patterns. How distributed white matter trajectory systems relate to one another therefore remains comparatively underexplored.

SILICA (Streamline Independent Component Analysis) offers a different way to organize the tractogram [5]. By decomposing streamline spatial fingerprints, SILICA identifies coherent, data-driven white-matter systems without first defining regional nodes. Critically, these systems are not detached latent factors, instead each remains explicitly linked to the streamlines through which it is expressed. SILICA therefore provides a representation in which tractography can be viewed not only as a collection of individual trajectories, but also as a set of trajectory-resolved structural systems.

This raises a new question, i.e., is there a connectome hidden among the SILICA components themselves? Here, we introduce Structural Network Coupling (SNC) to address this question. SNC uses SILICA components as the units of network organization and characterizes their relationships through patterns of expression across the tractogram. Rather than defining structural organization solely through direct connections between spatial nodes, SNC captures coupling among trajectory-resolved white-matter systems while preserving a direct route back to the trajectories that define them. It therefore provides an alternative structural connectivity representation in which network edges reflect relationships between distributed structural components rather than direct anatomical linkage alone. This creates a new network-level view of tractography and opens a natural path toward graph-based analysis and future structural– functional integration.

## 2. STRUCTURAL NETWORK COUPLING

### 2.1. SILICA Representation

SILICA represents whole-brain tractography through a sparse streamline-by-voxel fingerprint matrix *M*∈*R*^*N*streamline×*N*voxel^, and composes this representation as

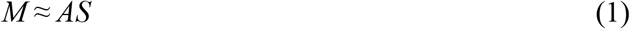

where *A*∈*R*^*N*streamline×*K*^ contains the streamline-loading profiles of *K* structural components, and *S*∈*R* ^*K*×*N*voxel^ contains their corresponding spatial sources.

A defining property of SILICA is that each component remains directly linked to the original tractography. The loading profile *A*(:,*k*) assigns a weight to every streamline for component *k*, allowing the component to be visualized directly as a component-weighted tractogram. Hence, SILICA represents each structural component simultaneously in three domains: spatial source maps, streamline loadings, and white matter trajectories.

While SILICA identifies the structural components themselves, the relationships among their streamline-loading profiles provide an additional level of organization.

### 2.2. From SILICA Loadings to Network Coupling

For subject *s*, SILICA provides a subject-specific loading matrix *A*_*s*_∈*R*^*N*streamline×*K*^. Each column *a*_*s,i*_ = *A*_*s*_(:,*i*) describes how component *i* is expressed across the subject’s trajectory population.

We define Structural Network Coupling (SNC) as the relationship between pairs of SILICA streamline-loading profiles:

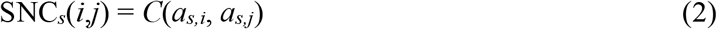

where *C* denotes a coupling operator.

The resulting matrix SNC_*s*_∈*R*^*K*×*K*^ provides a compact subject-level representation of relationships among trajectory-resolved structural components.

Conceptually, *A*_*s*_ → SNC_*s*_, transforming a streamline-by-component representation into a component-by-component structural network.

### 2.3. A Structural Coupling Representation

For the present analysis, *C* was defined as cosine similarity, equivalent to uncenter correlation, between the absolute streamline-loading profiles of each component pairs. It yields a cosine-normalized magnitude coupling:

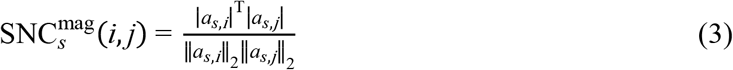

This measure satisfies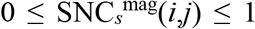, with higher values indicate that two components are strongly expressed across similar portions of the streamline population, whereas lower values indicate more distinct trajectory participation. Using loading magnitude makes the representation insensitive to the arbitrary sign orientation of ICA components.

Importantly, the general SNC framework is

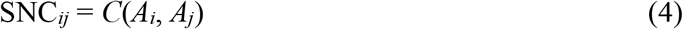

rather than any single choice of *C*.

Alternative formulations may retain complementary information, including cosine coupling,

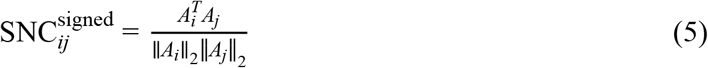

or centered correlation coupling,

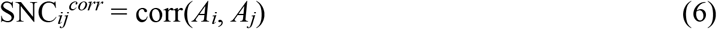

This flexibility allows the coupling representation to be adapted to different structural or multimodal questions without changing the underlying SILICA decomposition.

### 2.4. Why SILICA Makes This Possible

An SNC node is not an abstract graph node. Each node corresponds to a SILICA structural component and can be mapped directly back to its component-weighted tractogram.

Thus, every SNC relationship has three simultaneous representations: network relationship ↔ structural components ↔ white-matter trajectories.

This direct correspondence is a defining feature of SILICA-SNC: network organization remains linked to the trajectory anatomy from which it was derived.

### 2.5. A Different Level of the Structural Connectome

SNC is not intended to replace conventional structural connectivity. Rather, the two representations describe different organizational levels.

In conventional structural connectivity, node = anatomical regions, and edge = connection strength between regions.

In SNC, node = SILICA structural component, and edge = coupling between streamline-loading profiles.

Thus,

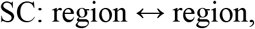

whereas

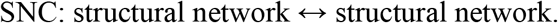

SNC therefore provides a component-level view of structural organization while retaining direct access to the trajectories defining each network node.

### 2.6. Toward Functional Network Integration

The SNC formulation also creates a natural conceptual parallel with ICA-based functional network connectivity.

Functional ICA produces spatial components together with their associated time courses, from which

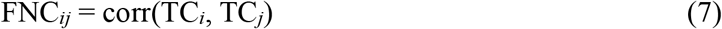

is commonly constructed [6].

SILICA similarly produces spatial structural components together with their streamline-loading profiles. Using the SNC definition in (6), this yields the parallel mappings

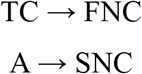

where FNC captures coupling between temporal loadings, whereas SNC captures coupling between structural components through their streamline-loading profiles.

This coupling operator most suitable for future structural-functional integration is intentionally left open. The present work focuses on establishing SNC as a structural network representation while preserving flexibility for future joint FNC-SNC analysis.

## 3. DEMONSTRATION

### 3.1. From Whole-Brain Tractography to SNC

SNC was demonstrated using diffusion MRI and whole-brain tractography from 30 healthy participants previously analyzed with SILICA [5]. Diffusion data were acquired at 2mm isotropic resolution using a conventional single-shell DTI protocol. Whole-brain tractography was reconstructed from the tensor model using an Euler integration scheme, with trajectories sampled at 0.5mm intervals.

Tractograms were transformed into MNI space and converted into the sparse streamline-by-voxel representation used by the SILICA framework. Group SILICA estimated 50 independent structural components, of which 41 were retained following visual assessment as anatomically interpretable white-matter components. The retained components captured coherent trajectory patterns spanning commissural, projection, and association systems and could be visualized directly as component-weighted tractograms.

Subject-specific back-reconstruction provided a streamline-loading profile for each retained component. SNC was then constructed independently for each participant from pairwise cosine similarity between absolute streamline-loading profiles, yielding a 41×41 subject-specific structural coupling matrix. Each node therefore represented a trajectory-resolved SILICA component, while each edge quantified shared streamline-loading structure between a pair of components.

Fig. 1 illustrates this progression in an example subject, from whole-brain tractography, through SILICA decomposition and component-weighted tractograms, to the resulting subject-specific SNC matrix. These individual SNC matrices formed the basis for the subsequent group-level analyses.

**Figure 1.**
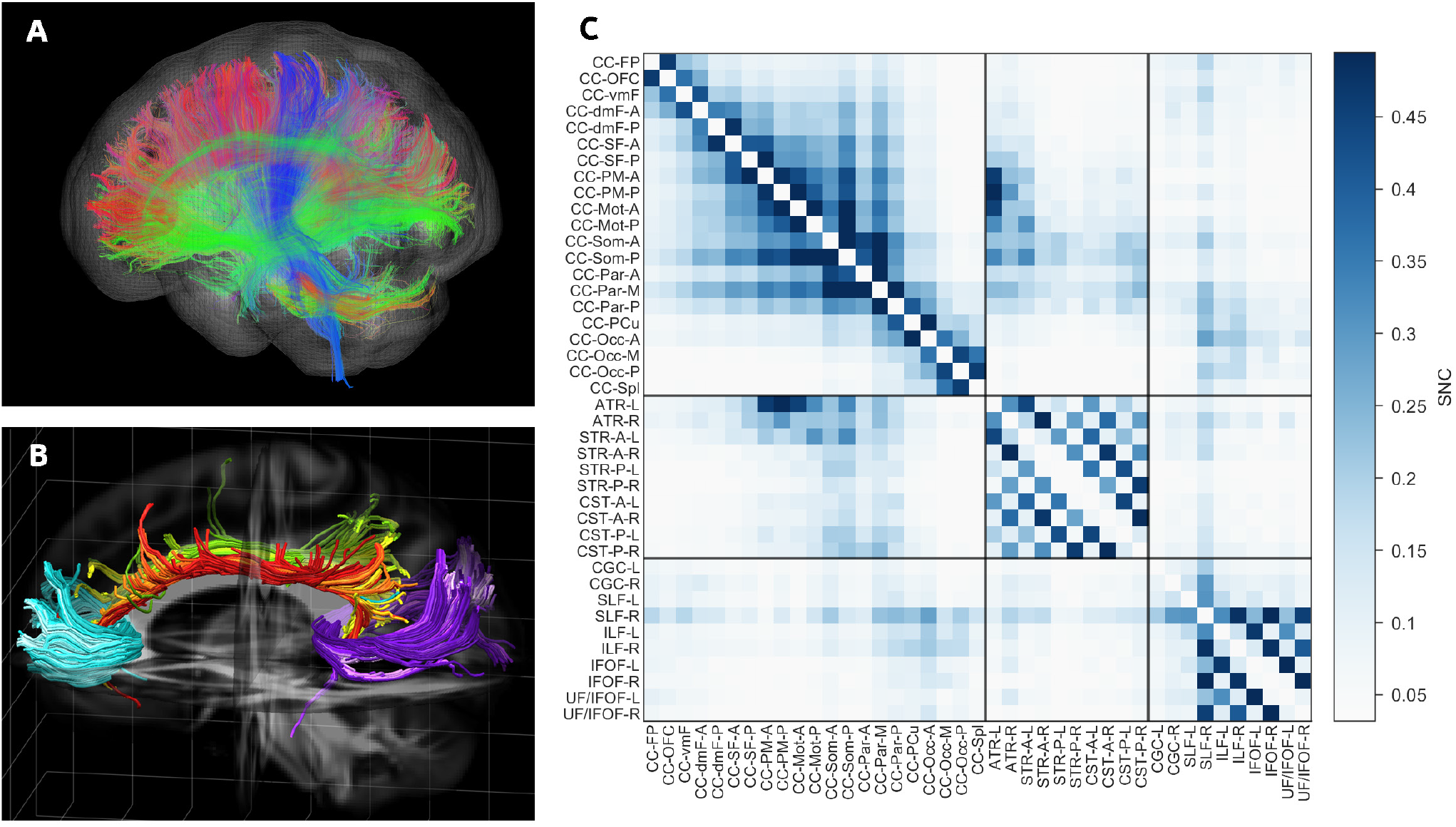
From Whole-Brain Trajectories to SNC. (A) Whole-brain tractography from an example subject. (B) Representative component-weighted tractograms obtained from SILICA back-reconstruction in the same subject. (C) Corresponding subject-specific SNC matrix, illustrating the transformation from trajectory-resolved structural components to a weighted structural network.

### 3.2. SNC Reveals Shared Trajectory Structure

To examine the anatomical meaning of individual SNC edges, representative high-, intermediate-, and low-coupling relationships were selected from the group-average SNC distribution and visualized using component-weighted tractograms from the same example subject shown in Fig. 1, avoiding the need to construct a pseudo group-average tractogram.

Using the right IFOF component as a common reference, coupling with the right UF/IFOF component was strong (0.564 ± 0.074), coupling with the left ILF component was intermediate (0.104 ± 0.021), and coupling with the left posterior superior thalamic radiation was weak (0.021 ± 0.008). The corresponding SNC values in the example subject were 0.538, 0.095, and 0.018, respectively (Fig. 2).

**Figure 2.**
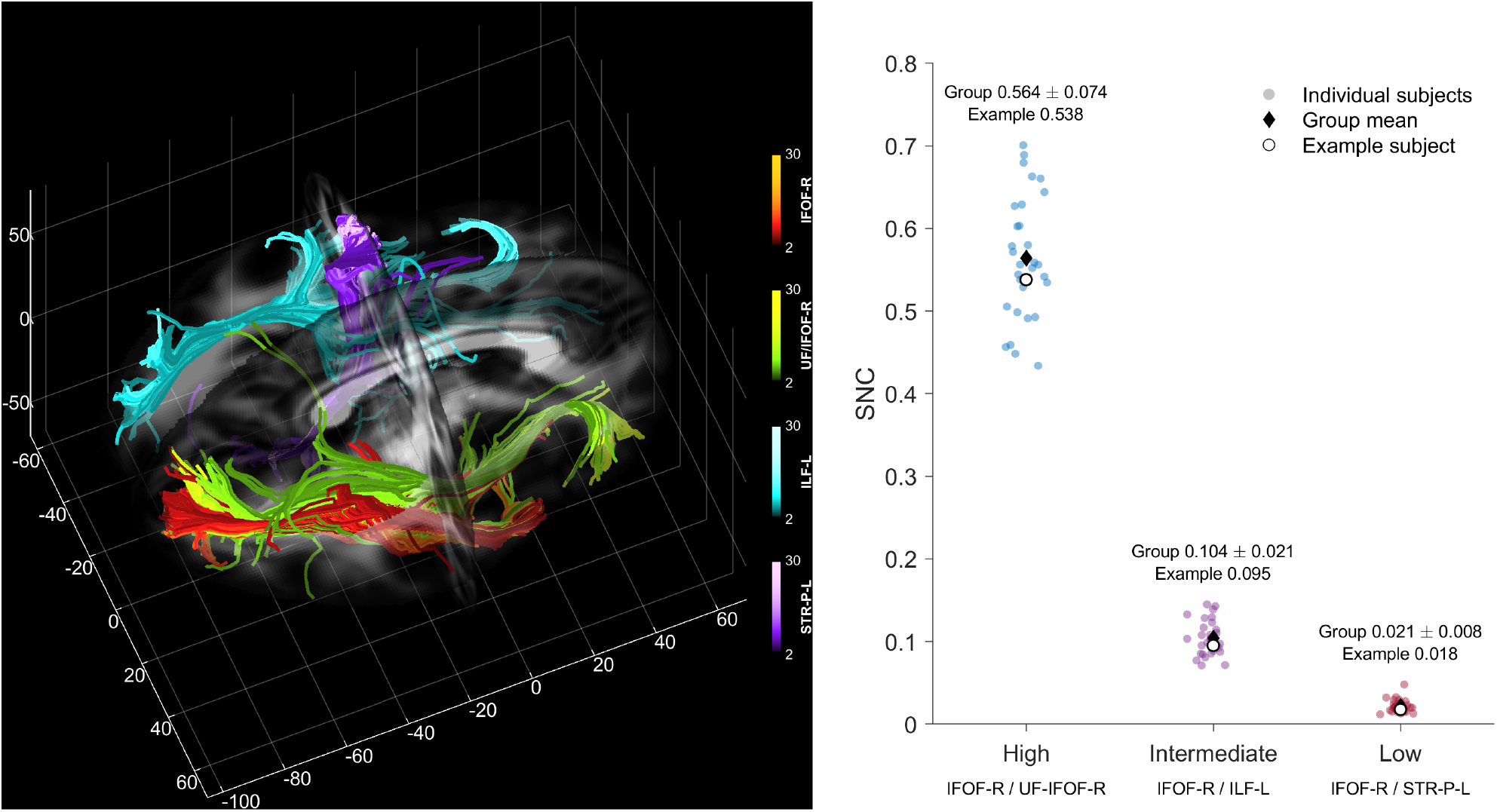
Anatomical interpretation of SNC edge strength. Representative high-, intermediate-, and low-coupling relationships identified from the group SNC are shown together with component-weighted tractograms from the example subject. Subject-level SNC distributions across the 30 participants demonstrate the consistency of the three coupling regimes, while the anatomical renderings illustrate the corresponding degree of shared trajectory structure.

The graded coupling strengths were consistent with the degree of shared trajectory structure. The strongly coupled pair comprised overlapping right-sided ventral association pathways, whereas the intermediate relationship involved anatomically related but hemispherically separated ventral association components. The weakly coupled pair belonged to different pathway classes and hemispheres and showed little shared streamline participation.

Because SILICA components are estimated in a data-driven manner rather than constrained to atlas-defined bundles, individual components may contain portions of neighboring anatomical pathways. Accordingly, SNC should not be interpreted as a measure of tract-name correspondence, but rather as a measure of cosine-normalized magnitude coupling between component-specific streamline-loading profiles. Higher SNC therefore indicates that the same portions of the streamline population carry large loading magnitudes for both components, whereas lower SNC indicates more distinct trajectory participation.

### 3.3. From SNC to Structural Network

Subject-specific SNC matrices were averaged across the 30 participants to characterize the group structural coupling architecture. The resulting matrix showed pronounced organization across the 41 structural components. To examine this structure without imposing anatomical class information, the group SNC was converted to a dissimilarity representation and subjected to agglomerative hierarchical clustering with optimal leaf ordering.

A striking correspondence emerged between the unsupervised SNC hierarchy and the anatomical organization previously assigned from visual inspection of the SILICA components [5]. Importantly, the commissural, projection, and association labels had been assigned from component anatomy and were not used in either SNC construction or hierarchical clustering. Nevertheless, reordering the same group SNC from its original anatomical arrangement (Fig. 3A) according to the data-driven hierarchy (Fig. 3B–C) separated the components into three fully contiguous systems, projection, commissural, and association pathways, with only two class transitions, the theoretical minimum.

**Figure 3.**
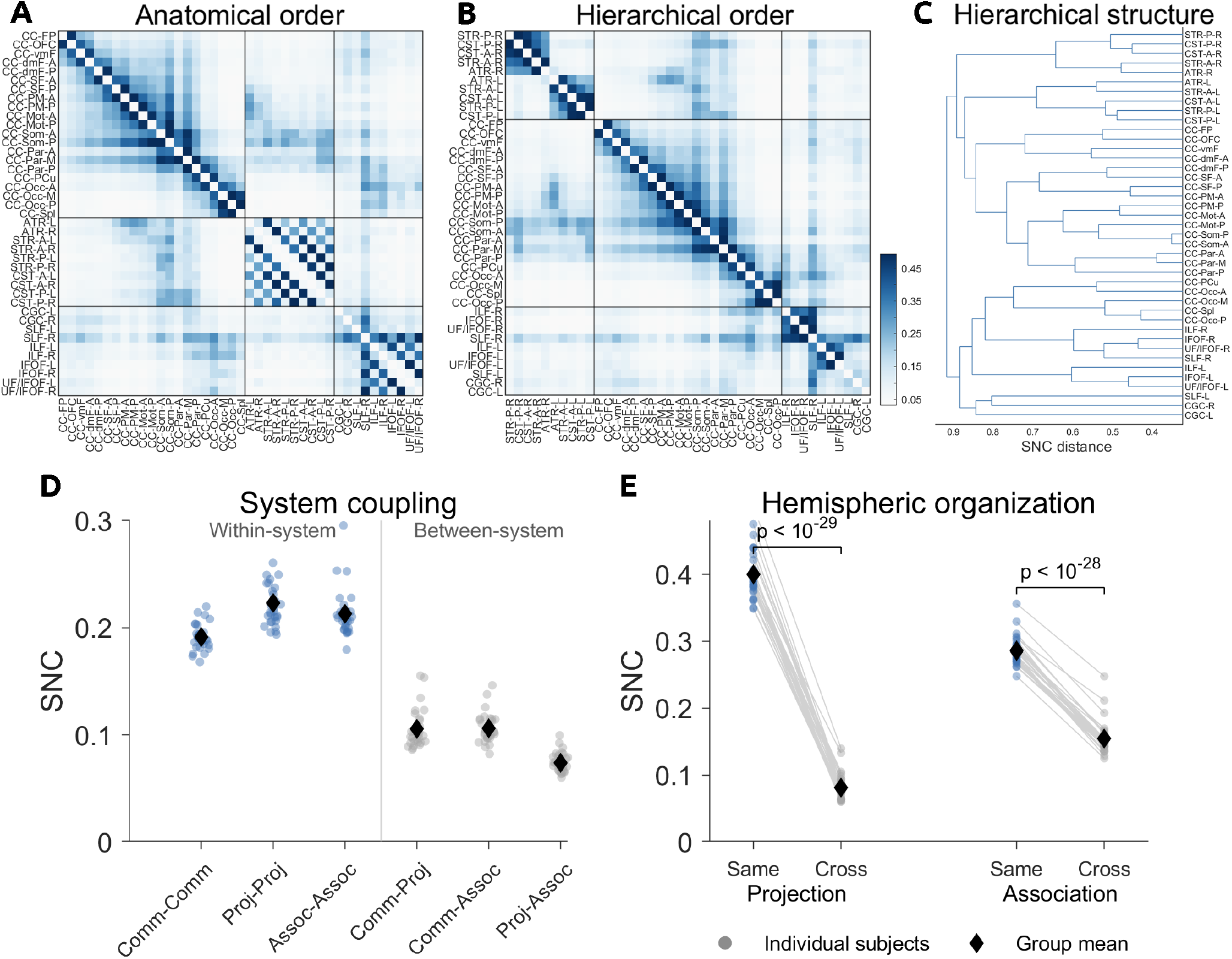
Hierarchical organization of group SNC. (A) Group SNC in the original anatomical order. (B) Hierarchically reordered SNC. (C) Corresponding dendrogram. (D) Within-and between-system coupling. (E) Same-versus cross-hemisphere coupling in projection and association systems.

In the original anatomical ordering, left/right homologues were listed adjacently, producing a characteristic checkerboard pattern in the unilateral projection and association systems (Fig. 3A). In contrast, the data-driven hierarchical ordering grouped components predominantly by hemisphere, yielding contiguous right-and left-sided blocks within these systems (Fig. 3B–C).

Random permutation of the anatomical class labels never produced an equally contiguous ordering in 100000 permutations (*p* < 10^−5^). The exact probability of obtaining three perfectly contiguous groups with the observed class sizes by random ordering was 1.21 × 10^−16^.

The hierarchy also recovered organization below the level of the three major tract classes. Projection components separated almost completely by hemisphere, with the five right-sided components followed by the five left-sided components. Association pathways showed a similar hemispheric organization, with right-and left-sided association components forming largely contiguous subgroups and the bilateral cingulum pair occupying a distinct terminal position. Commissural components formed a continuous sequence within the central portion of the hierarchy, consistent with their graded anterior-to-posterior callosal organization.

Quantitative analysis confirmed the system-level block structure observed in the hierarchically ordered SNC (Fig. 3D). Mean within-system coupling was 0.191 ± 0.012 for commissural pathways, 0.223 ± 0.023 for projection pathways, and 0.213 ± 0.022 for association pathways. Cross-system coupling between commissural and projection components (0.105 ± 0.018) and between commissural and association components (0.106 ± 0.014) was greater than direct projection–association coupling (0.074 ± 0.009, *p* = 1.7 × 10^−11^ and 1.2 × 10^−15^, respectively).

A particularly strong hemispheric effect was observed within the unilateral tract systems (Fig. 3E). Projection components showed substantially greater same-hemisphere than cross-hemisphere coupling (0.400 ± 0.036 versus 0.081 ± 0.020, *p* = 2.1 × 10^−30^). Association components showed the same pattern (0.286 ± 0.021 versus 0.154 ± 0.025, *p* = 9.9 × 10^−29^).

These findings indicate that SNC contains organization at multiple spatial scales: broad separation among major white-matter systems, strong hemispheric segregation within unilateral pathways, and finer coupling structure within each tract family. The relatively stronger coupling of commissural components with both projection and association systems, compared with the weaker direct projection–association coupling, is also consistent with the cross-hemispheric and cross-system anatomical role of the commissural pathways. This pattern should be interpreted as a structural coupling relationship rather than evidence of causal communication.

Notably, hierarchical reordering does not alter the SNC values or impose the observed blocks, instead it only changes node order according to similarity in coupling profiles. The close agreement between the independently assigned anatomical classes and the data-driven SNC hierarchy therefore provides convergent evidence that SNC preserves meaningful white-matter organization in network space.

### 3.4. Network Consistency

The stability of the group SNC architecture was evaluated using repeated random split-half analysis. In each of 1,000 iterations, the 30 participants were randomly divided into two groups of 15, separate group SNC matrices were estimated, and similarity between their unique edge profiles was quantified using Pearson correlation.

Split-half SNC correlations were consistently high and narrowly distributed (mean *r* = 0.993, median *r* = 0.993, 95% empirical interval 0.988-0.996, range 0.985-0.996, Fig. 4). These results indicate that the group-level SNC architecture is highly stable across different participant subsets.

**Figure 4.**
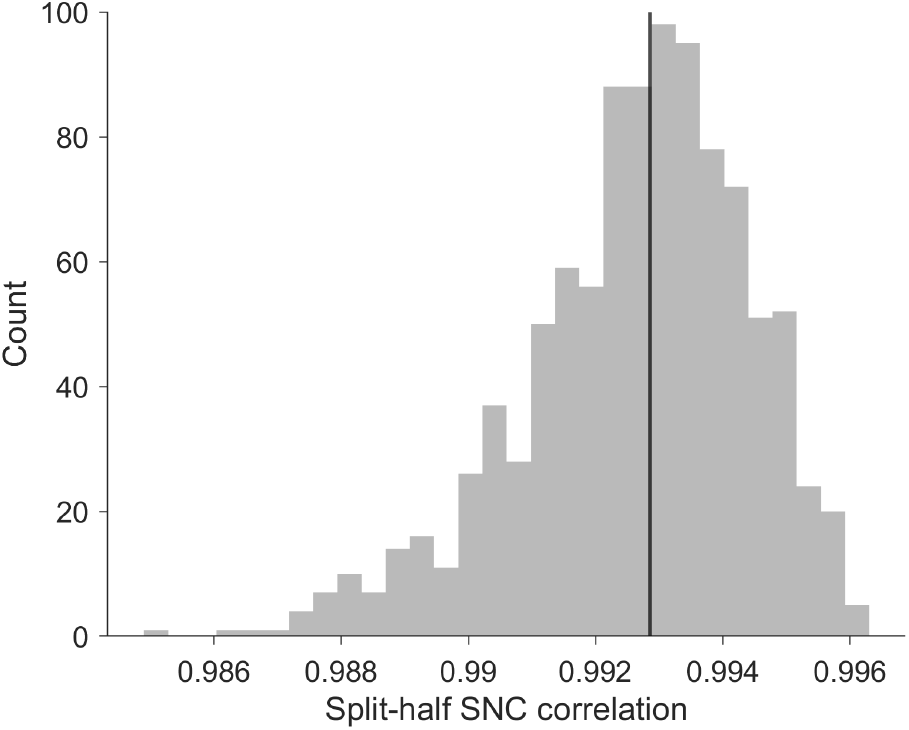
Split-half consistency of group SNC. Distribution of Pearson correlations between group SNC edge profiles across 1,000 random 15/15 participant splits.

These results indicate that the group SNC structure is not driven by a particular subset of participants and provide an initial demonstration of reproducibility of the trajectory-based network representation.

## 4. CONCLUSION

SILICA provides a trajectory-resolved decomposition of whole-brain tractography. SNC extends that representation by characterizing relationships among SILICA components through their streamline-loading profiles.

The conceptual progression is simple:

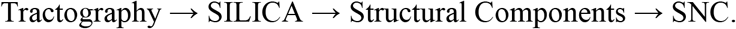

Unlike conventional region-based connectomes, SNC uses data-driven white-matter components as network nodes while preserving direct access to the trajectories defining each component.

The formulation SNC_*ij*_ = *C*(*A*_*i*_, *A*_*j*_) also keeps the network representation flexible, allowing different coupling measures to emphasize shared trajectory participation, signed relationships, or future multimodal correspondence.

SILICA decomposes the tractogram. SNC connects the components. Together, they provide a direct route from trajectory-scale white-matter organization to network-scale structural representation and establish a foundation for future structural-functional network integration.

## ACKNOWLEDGMENT

This work was supported by National Institutes of Health grants 1R01EB006841, 1R01EB005846, R01AG090597 and National Science Foundation grant 2112455. The authors declare no conflicts of interest relevant to the content of this manuscript.

## COMPLIANCE WITH ETHICAL STANDARDS

All subjects provided written informed consent, and study procedures were approved by the Institutional Review Board at the University of New Mexico / Mind Research Network (MRN), under the COBRE program.

